# Sensory-sleep dysfunction is a shared phenotype across genetically distinct neurodevelopmental disorder models

**DOI:** 10.64898/2026.08.31.748066

**Authors:** Jadwiga Bilchak, Kyla Mace, Jenny Luong, Kevin Wu, Matthew S. Kayser

## Abstract

Sensory abnormalities and sleep disruption frequently co-occur in neurodevelopmental disorders (NDDs), but how sensory input interacts with sleep regulation in NDDs remains poorly understood. Here, we examined vibration-induced sleep (VIS), in which prolonged gentle vibration promotes sleep in *Drosophila*, in three genetically distinct NDD models: *dNf1*, *dNrx1*, and *dFmr1*. Despite markedly different baseline sleep phenotypes, all three mutants exhibited impaired VIS, identifying disrupted sensory-sleep integration as a shared phenotype across these NDD models. Behavioral responses to vibration, activity-associated CRTC signaling in Nanchung-positive (Nan+) mechanosensory neurons, and thermogenetic activation of Nan+ neurons revealed distinct underlying abnormalities across the three models, indicating that disruption at different points along the sensory-sleep axis can converge on the same behavioral phenotype. The effect of mechanosensory stimulation was also strongly shaped by homeostatic sleep drive: following sleep deprivation, vibration promoted sleep in *dNf1* mutants, remained ineffective in *dNrx1* mutants, and opposed recovery sleep in *dFmr1* mutants. Together, these findings identify impaired sensory regulation of sleep as a point of convergence across genetically distinct NDD models and demonstrate that the expression of this shared phenotype depends on internal state.

## INTRODUCTION

Sensory information must be continuously evaluated to determine whether it warrants arousal or can be safely ignored. This transformation is not fixed: the behavioral effect of a stimulus depends on its intensity, salience, and temporal structure, as well as the animal’s internal state. Even during sleep, animals retain the ability to discriminate among sensory signals, and the probability that a stimulus produces awakening can be modulated by physiological state (1–5). Conversely, certain repetitive sensory stimuli can facilitate rather than disrupt sleep. In humans, gentle rocking promotes sleep onset and alters sleep-associated neural oscillations, whereas low-intensity vibration can reduce wakefulness after sleep onset and improve subjective sleep quality (6,7). The sleep-promoting effects of rhythmic mechanosensory stimulation are conserved across species. Rocking increases non-rapid eye movement sleep and accelerates sleep onset in mice through vestibular input (8), while prolonged gentle vibration promotes sleep in *Drosophila melanogaster* (*9,10*). Vibration-induced sleep (VIS) becomes more pronounced with repeated stimulation and provides cognitive and physiological benefits, supporting the conclusion that it represents a functional sleep state rather than behavioral immobility alone (11).

The interaction between sensory processing and sleep may be particularly important in neurodevelopmental disorders (NDDs), as sensory over-responsivity is associated with sleep disturbances in autistic children (12–14). More broadly, sleep disturbances are common across NDDs, but their expression is highly heterogeneous, including differences in sleep duration, timing, continuity, and other aspects of sleep architecture (15–18). This heterogeneity is also evident in mammalian models, in which different NDD-associated mutations preferentially disrupt sleep continuity, sleep initiation, or sleep amount (19–22).

Sensory-processing abnormalities are likewise common and heterogeneous, encompassing hyperresponsiveness, hyporesponsiveness, sensory seeking, and sensory avoidance across multiple modalities and varying substantially even among individuals with the same genetic diagnosis (23–25). Although sleep and sensory phenotypes are each well recognized in NDDs, they are typically considered as separate features. Whether sensory regulation of sleep is itself disrupted in NDDs remains poorly understood.

VIS in *Drosophila* provides a tractable system for investigating how sensory input influences sleep. Gentle vibration is detected by peripheral mechanosensory neurons expressing the TRPV channel Nanchung (Nan), including Johnston’s organ neurons in the antennae, and activation of Nan+ neurons is sufficient to promote sleep in the absence of vibration (9,10). Conversely, disrupting antennal mechanosensory input reduces VIS, establishing Nan+ sensory neurons as an important entry point through which mechanosensory stimulation influences sleep. The ability to examine sensory responsiveness, Nan+ neuron signaling, and the sleep-promoting effects of Nan+ activation enables sensory-sleep dysfunction to be assessed at multiple levels in NDD models.

Here, we examined VIS in three *Drosophila* models of NDDs with mutations in *Nf1*, *Neurexin*, or *Fmr1*. Despite distinct baseline sleep phenotypes, all three mutants exhibited markedly impaired VIS. Behavioral responses to vibration, activity-associated CRTC signaling in Nan+ sensory neurons, and thermogenetic activation of Nan+ neurons revealed different abnormalities across the three models, indicating that disruption at distinct points along the sensory-sleep axis can converge on a common impairment in sensory-induced sleep. Finally, manipulating homeostatic sleep pressure revealed that the effect of mechanosensory stimulation is strongly state dependent, promoting sleep in sleep-deprived *dNf1* mutants, remaining ineffective in *dNrx1* mutants, and opposing recovery sleep in *dFmr1* mutants. Together, these findings identify impaired sensory-sleep integration as a shared phenotype across genetically distinct NDD models and demonstrate that its expression depends on both the underlying mutation and internal sleep state.

## RESULTS

### Vibration-induced sleep is impaired in multiple *Drosophila* models of neurodevelopmental disorders

To investigate sensory-sleep integration in neurodevelopmental disorders (NDDs), we used vibration-induced sleep (VIS), a behavioral paradigm in which prolonged gentle vibration promotes daytime sleep in *Drosophila* (9–11). We examined flies carrying mutations in *Nf1* (*dNf1*), *Neurexin (dNrx1),* or *Fmr1* (*dFmr1*), together with their respective genetic controls. Adult female flies were first monitored for 24 h under standard conditions to establish baseline sleep. Beginning at ZT0 (lights on), flies were exposed to continuous gentle vibration, and VIS was quantified during the 12-h daytime period as the change in sleep relative to the corresponding baseline day (Fig. 1A).

**Figure 1:**
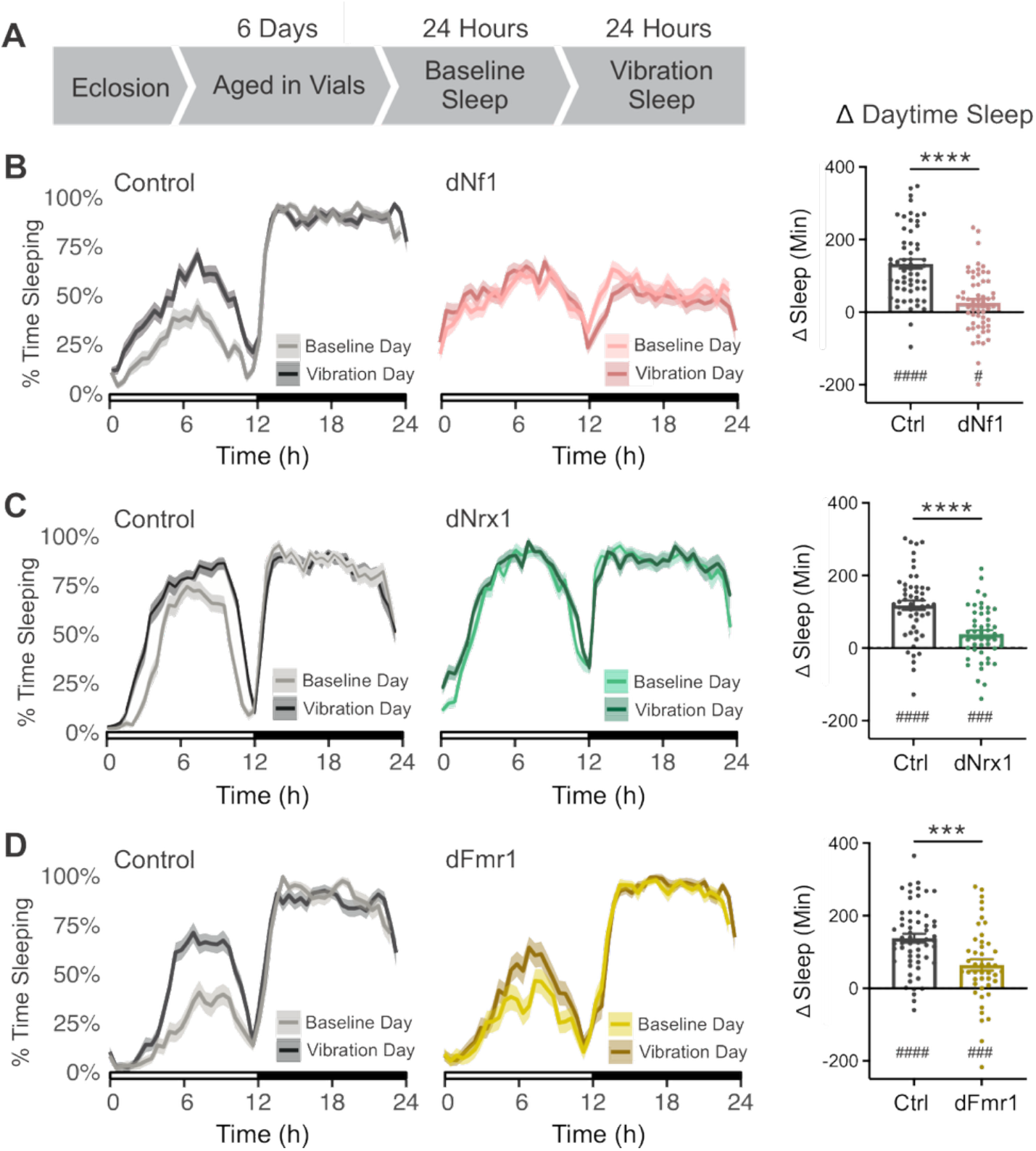
Vibration-induced sleep is impaired in multiple *Drosophila* models of neurodevelopmental disorders. **(A)** Experimental design. Following eclosion, female flies were aged in vials for 6 days. Sleep was then recorded for 24 h under baseline conditions, followed by 24 h of continuous gentle vibration beginning at ZT0. **(B–D)** Averaged 24-h sleep traces during the baseline and vibration days for: *dNf1* mutants and their corresponding genetic control (**B**), *dNrx1* mutants and control (**C**), and *dFmr1* mutants and control (**D**). Lines show the mean and shaded regions indicate SEM. Right graphs show vibration-induced sleep, calculated for each fly as the change in daytime sleep between the vibration and baseline days (Δ daytime sleep = vibration-day sleep − baseline-day sleep, ZT0–12). Gentle vibration produced a robust increase in daytime sleep in each control group. In contrast, *dNf1* and *dNrx1* mutants showed little or no increase in daytime sleep, whereas *dFmr1* mutants exhibited a smaller increase than their corresponding controls. Points represent individual flies; bars show mean ± SEM. **(B)** Control, *n* = 59; *dNf1*, *n* = 58. **(C)** Control, *n* = 52; *dNrx1*, *n* = 49. **(D)** Control, *n* = 58; *dFmr1*, *n* = 45. Changes in daytime sleep were compared between genotypes using unpaired t-tests. \*\*\**P* < 0.001; \*\*\*\**P* < 0.0001. Changes in daytime sleep were compared within genotypes to a hypothetical mean of zero using one-sample t-tests ^#^*P<0.05*; ^###^*P<0.001*; ^####^*P<0.0001*

Consistent with previous reports, the three NDD models exhibited distinct baseline sleep phenotypes (Fig. 1B-D; Fig. S1). dNf1 mutants displayed severe sleep fragmentation and disrupted circadian rhythms (26,27), dNrx1 mutants showed milder sleep fragmentation (28), whereas baseline sleep in dFmr1 mutants was largely comparable to that of genetic controls. Despite these markedly different baseline phenotypes, all three mutants exhibited impaired VIS. Gentle vibration robustly increased daytime sleep in each control strain, whereas VIS was nearly absent in dNf1 and dNrx1 mutants and markedly attenuated in dFmr1 mutants (Fig. 1B-D).

Because baseline sleep differed substantially among the three NDD models, we next asked whether these differences could account for the reduced VIS observed in mutant flies (Fig. S2). We compared vibration-day sleep between mutants and controls while accounting for each fly’s baseline sleep. For dNf1 and dFmr1, genotype remained significantly associated with vibration-day sleep after accounting for baseline sleep, indicating that impaired VIS is not explained by baseline sleep differences alone. In contrast, the effect of dNrx1 genotype was no longer significant after accounting for baseline sleep, suggesting that the elevated baseline sleep of dNrx1 mutants may contribute to reduced magnitude of VIS. High baseline sleep alone, however, was not sufficient to prevent VIS. Male flies normally sleep more than females during the day (29,30), so we examined VIS in male mutants and controls. Control males exhibited high levels of baseline daytime sleep, significantly higher than their Nrx1 mutant counterparts (Fig. S3 B), and comparable to that observed in female dNrx1 mutants, yet showed a robust increase in sleep during vibration (Fig. S3). Similar genotype-dependent reductions in VIS were also observed in males, indicating that impaired VIS is not sex-specific. Because higher baseline daytime sleep in males reduced the dynamic range for detecting further increases in sleep, females were used for subsequent experiments. Thus, despite substantial differences in baseline sleep architecture, loss of dNf1, dNrx1, or dFmr1 converges on a reduced ability of mechanosensory stimulation to promote sleep.

### Impaired VIS is not explained by a common deficit in vibration sensitivity

We next asked whether impaired VIS in the NDD mutants resulted from altered sensitivity to vibration. Control flies exhibit a transient increase in locomotor activity at vibration onset, providing a behavioral measure of vibration detection (9). We therefore recorded locomotor activity for 30 min before vibration onset and throughout a subsequent 30 min vibration period. To avoid conflating sleep-wake transitions with vibration-evoked activity, flies that were asleep when vibration began were excluded from analysis. As expected, control flies showed a pronounced increase in activity during the first 5 min of vibration (Fig. 2 A, C). dNf1 mutants exhibited a comparable onset response, demonstrating that they appropriately detect the vibration stimulus under these conditions (Fig. 2 B). By contrast, dNrx1 mutants did not increase their activity at vibration onset (Fig. 2 D). dFmr1 mutants displayed elevated activity during the pre-vibration period in this assay and showed no further increase after vibration began (Fig. 2 E), complicating interpretation of the population-level vibration response.

**Figure 2:**
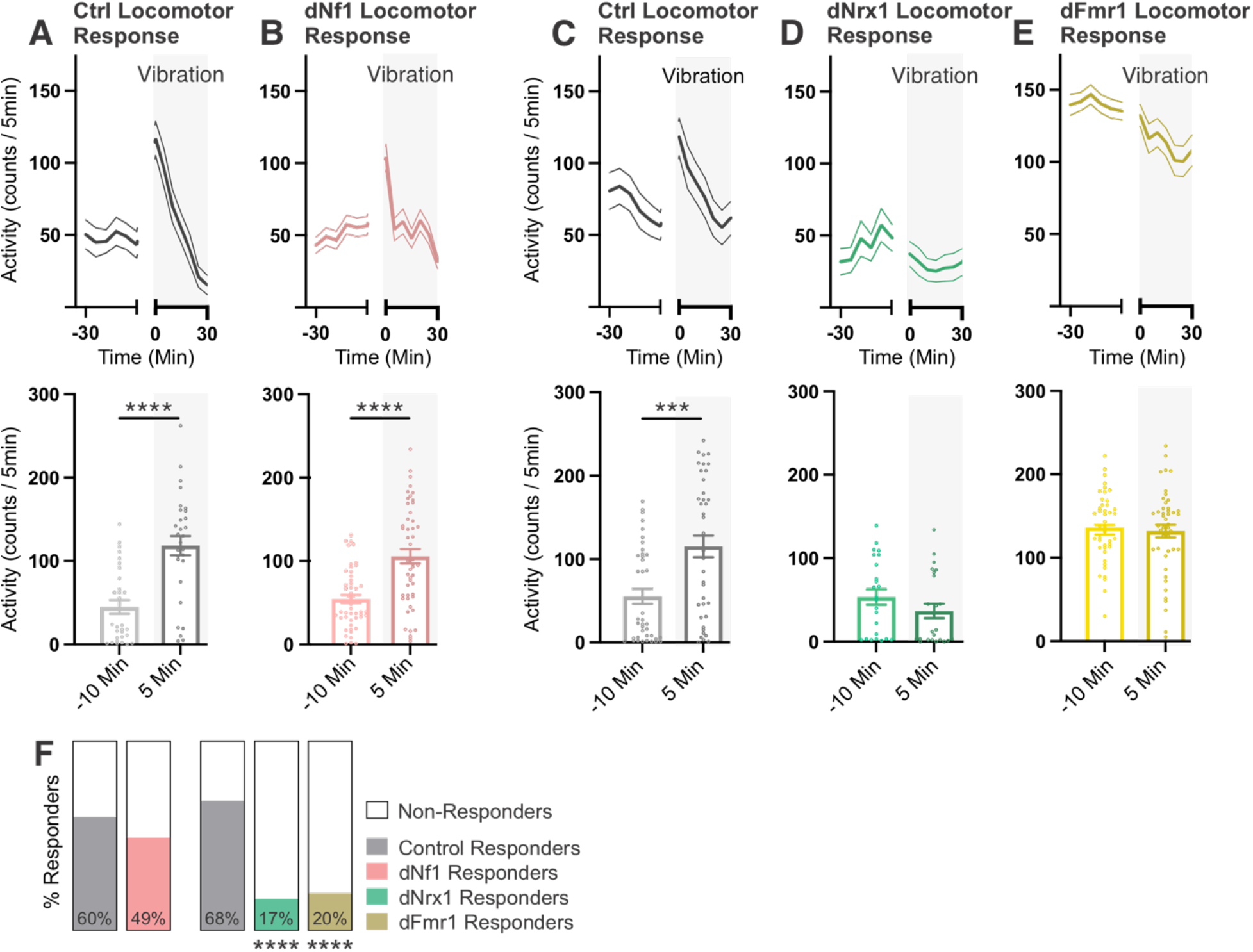
Vibration-evoked locomotor responses are reduced in *dNrx1* and *dFmr1* mutants. Locomotor activity was recorded before and during a 30 min period of continuous gentle vibration beginning at ZT3. Only flies that were awake at vibration onset were included in the analysis. Activity counts were collected in 1-min bins and averaged into 5-min bins. **(A–E)** Locomotor activity in the genetic control for *dNf1*, n = 30 (**A**), *dNf1* mutants, n = 49 (**B**), the shared genetic control for *dNrx1* and *dFmr1*, n = 39 (**C**), *dNrx1* mutants, n = 24 (**D**), and *dFmr1* mutants, n = 46 (**E**). Upper graphs show activity over time; gray shading denotes the vibration period. Central lines represent the mean and flanking lines indicate SEM. Lower graphs show activity during selected 5-min intervals before vibration (−10 min), and within the first 5 minutes of vibration (5 min). Control flies exhibited a transient increase in locomotor activity at vibration onset. *dNf1* mutants displayed a similar vibration-onset response. In contrast, *dNrx1* mutants did not significantly increase activity at vibration onset. *dFmr1* mutants exhibited elevated baseline activity and no further increase at vibration onset. Points represent individual flies; bars show mean ± SEM. Activity before and after vibration was analyzed using a paired t-test. **(F)** Percentage of flies classified as responders to vibration onset. Flies were defined as responders when their absolute change in activity during the first 5 min of vibration exceeded two standard deviations of their activity during the 15 min immediately preceding vibration. The proportion of responders did not differ significantly between *dNf1* mutants (49%) and their genetic control (60%). In contrast, *dNrx1* (17%) and *dFmr1* (20%) mutants were significantly less likely to respond than their shared genetic control (68%). Responder proportions were compared using a Fisher’s exact test. \*\*\**P* < 0.001; \*\*\*\**P* < 0.0001.

Because differences in baseline activity could obscure vibration-evoked responses, we also classified individual flies according to the magnitude of their response relative to baseline activity. Flies were designated as responders if the absolute change in activity at vibration onset exceeded two standard deviations of their activity during the 15-min baseline period, thereby capturing both increases and decreases in locomotor activity. Consistent with the population-level measurements, the proportion of dNf1 mutants classified as responders did not differ significantly from controls (Fisher’s exact test, p=0.3588). In contrast, the proportion of responders was significantly reduced in both dNrx1 and dFmr1 mutants relative to their respective controls (p=0.0014 and p=0.0077, respectively; Fig. 2F).

To further characterize vibration responsiveness across genotypes, we used a custom-built vibration stage to expose flies to a range of stimulus intensities (Fig. S4). At the lowest intensity, few flies responded, and response rates were similar between mutants and controls. As stimulus intensity increased, response rates increased across genotypes, and at higher intensities all three mutant strains exhibited response rates comparable to their respective controls. Thus, despite differences in their responses at the standard VIS stimulus, all three mutants retained the capacity to behaviorally respond to stronger vibration.

Altered sensory processing could also manifest as hypersensitivity, raising the possibility that excessive mechanosensory input interferes with sleep induction. We therefore reduced mechanosensory input by surgically removing the antennae, which contain Johnston’s organ neurons that detect vibration and other forms of antennal displacement (31–33). Consistent with previous work (9), antennectomy reduced VIS in control flies (Fig. S5). In contrast, antennectomy did not restore VIS in any of the three mutants (Fig. S5), arguing against excessive antennal sensory input as a common cause of impaired VIS. Together, these results indicate that impaired VIS across the three NDD models is not explained by a common deficit in vibration sensitivity. Although acute behavioral responses to the standard VIS stimulus differed among genotypes, all three mutants retained the capacity to respond to stronger vibration, and reducing antennal sensory input did not restore VIS. Thus, distinct alterations in sensory processing may contribute to the phenotype in individual mutants, but do not provide a common explanation for their shared failure of vibration to promote sleep.

### Vibration evokes genotype-dependent changes in CRTC localization in Nan+ sensory neurons

Because behavioral responses to vibration may not directly reflect activity within the underlying mechanosensory neurons, we next examined vibration-evoked activity in nanchung-positive (Nan+) sensory neurons. We used CREB-regulated transcriptional coactivator (CRTC) as an activity-dependent reporter. CRTC undergoes rapid, activity-dependent nuclear translocation in *Drosophila* neurons, providing a bidirectional measure of recent neuronal activity (34). We expressed CRTC-GFP in Nan+ neurons and quantified its subcellular localization in Johnston’s organ neurons under baseline conditions and following vibration (Fig. 3 A).

**Figure 3:**
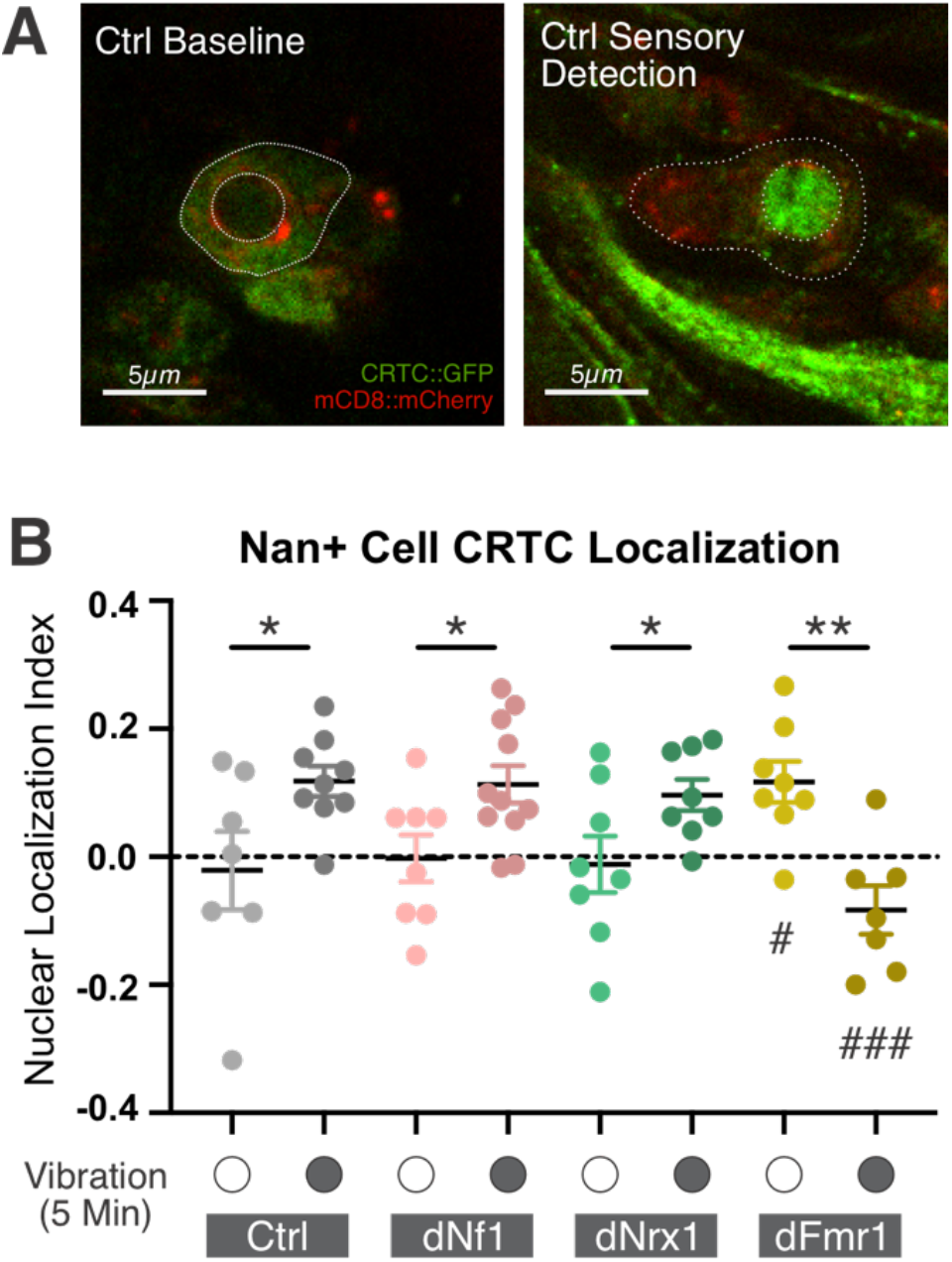
Vibration-evoked CRTC localization in Nan+ sensory neurons is altered in *dFmr1* mutants. **(A)** Representative confocal images of CRTC::GFP (green) and the membrane marker mCD8::mCherry (red) in Nan+ sensory neurons from control flies under sham conditions (left) or following 5 min of vibration (right). Dotted outlines indicate the nuclear and cytoplasmic regions of interest used to quantify CRTC localization. Scale bars, 5 µm. **(B)** Nuclear localization index (NLI) of CRTC::GFP in Nan+ sensory neurons from control, *dNf1*, *dNrx1*, and *dFmr1* flies under sham conditions or following 5 min of vibration. NLI was calculated as (mean nuclear GFP intensity − mean cytoplasmic GFP intensity)/(mean nuclear GFP intensity + mean cytoplasmic GFP intensity), with positive values indicating greater nuclear localization. Vibration significantly increased CRTC nuclear localization in control, *dNf1*, and *dNrx1* sensory neurons. In contrast, *dFmr1* neurons exhibited elevated CRTC nuclear localization under sham conditions and a significant reduction following vibration. Each point represents the mean NLI of five neurons sampled from one antenna of one fly. Bars show mean ± SEM. Sham: control, *n* = 7; *dNf1*, *n* = 8; *dNrx1*, *n* = 8; *dFmr1*, *n* = 8. Vibration: control, *n* = 9; *dNf1*, *n* = 11; *dNrx1*, *n* = 8; *dFmr1*, *n* = 7. Data were analyzed by two-way ANOVA with Holm–Šídák multiple-comparisons tests. Asterisks (*) indicate comparisons between sham and vibration within a genotype, whereas number signs (#) indicate comparisons with the control mean under the corresponding condition. \**P* < 0.05; \*\**P* < 0.01; #*P* < 0.05; ###*P* < 0.001.

We first examined CRTC localization following 5 min of vibration, corresponding to the period used to quantify acute behavioral responses to vibration. In control flies, vibration significantly increased nuclear CRTC localization in Nan+ neurons, consistent with activation of these mechanosensory neurons (Fig. 3 B). We next asked whether impaired VIS was associated with a failure to activate Nan+ sensory neurons. Despite reduced VIS, both dNf1 and dNrx1 mutants exhibited increased nuclear CRTC localization following vibration, indicating that Nan+ neurons remained responsive to the stimulus (Fig. 3 B). In contrast, dFmr1 mutants exhibited elevated nuclear CRTC signal at baseline that significantly decreased following vibration (Fig. 3 B).

These cellular responses only partially corresponded to the acute behavioral responses observed following vibration onset. dNf1 mutants exhibited both normal behavioral responsiveness and vibration-induced increases in nuclear CRTC, whereas dNrx1 mutants showed robust vibration-induced CRTC responses despite a reduced acute locomotor activation response. dFmr1 mutants also exhibited reduced acute behavioral responsiveness but showed a distinct cellular phenotype, with elevated nuclear CRTC signal at baseline that decreased following vibration. Thus, impaired VIS across these NDD models is not uniformly associated with a failure of vibration to activate Nan+ sensory neurons, indicating that the shared deficit in sensory-induced sleep is not due to a common defect at this early stage of sensory processing.

### Direct activation of Nan+ sensory neurons reveals distinct sites of sensory-sleep circuit dysfunction

We next asked whether direct activation of Nan+ sensory neurons could bypass mechanosensory stimulation and promote sleep in NDD mutants. To thermogenetically activate these neurons, we expressed the temperature-gated cation channel dTRPA1 under the control of *Nan-Gal4*. Sleep was recorded first at 22°C, below the activation threshold of dTRPA1, and subsequently at 29°C to activate dTRPA1-expressing neurons (Fig. 4A).

**Figure 4:**
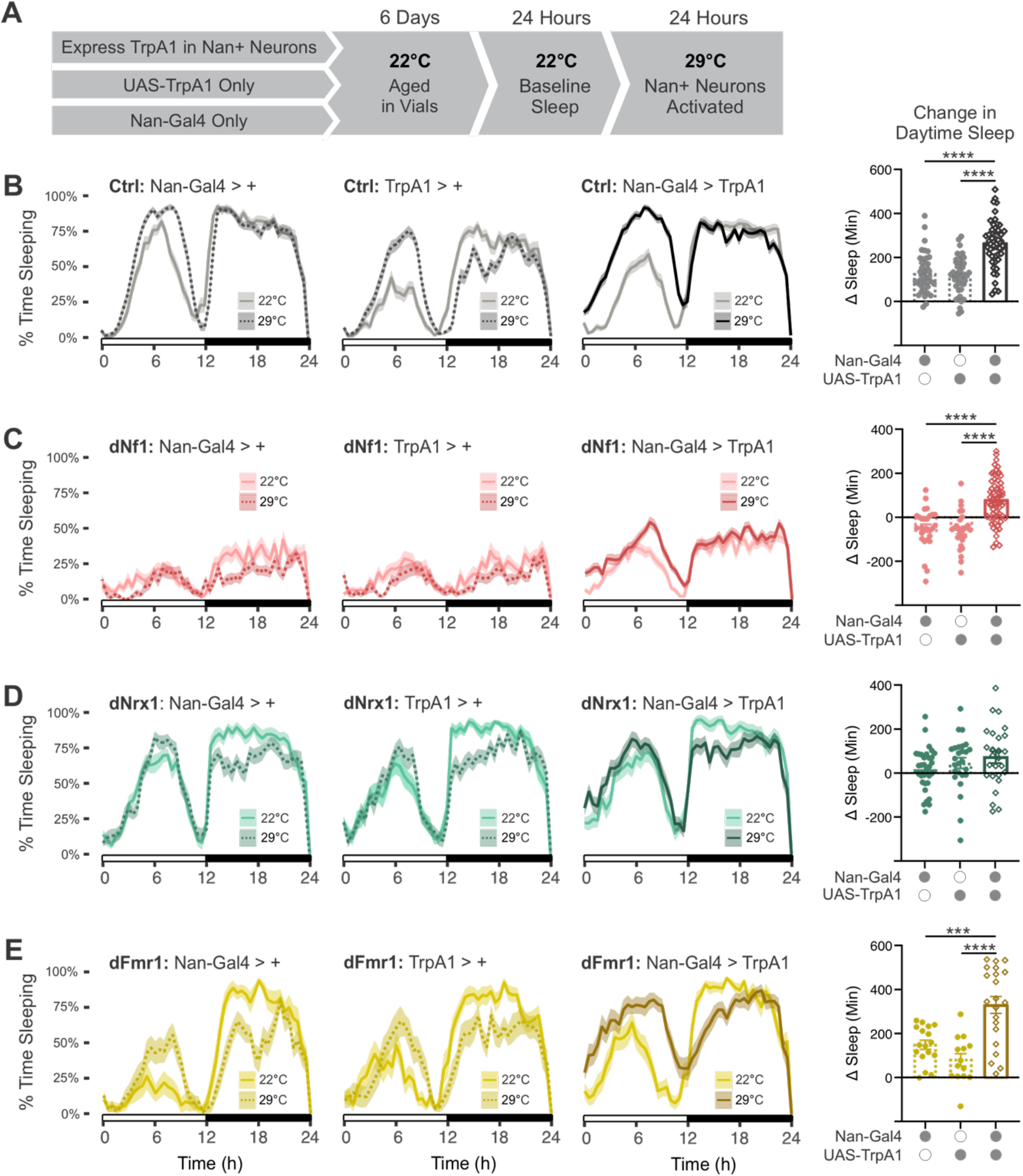
Direct activation of Nan+ sensory neurons differentially promotes sleep in NDD mutants. **(A)** Experimental design. Female flies expressing the temperature-sensitive cation channel TrpA1 in Nan+ sensory neurons (*Nan-GAL4 > UAS-TrpA1*) and their corresponding Gal4-only and UAS-only genetic controls were raised and aged for 6 days at 22°C. Sleep was recorded for 24 h at 22°C to establish baseline levels, followed by 24 h at 29°C to activate TrpA1-expressing neurons. **(B-E)** Averaged sleep traces at 22°C and 29°C and the corresponding change in daytime sleep for flies in control (**B**), *dNf1* (**C**), *dNrx1* (**D**), and *dFmr1* (**E**) genetic backgrounds. Within each background, the left and middle traces show the temperature response of the *Nan-GAL4/+* and *UAS-TrpA1/+* genetic controls, respectively, and the right trace shows flies expressing TrpA1 in Nan+ neurons. Right graphs show the change in daytime sleep following the temperature shift (Δ sleep = daytime sleep at 29°C − daytime sleep at 22°C). **(B)** In a control genetic background, activation of Nan+ neurons produced a robust increase in daytime sleep that was significantly greater than the temperature-dependent changes observed in both genetic controls. N = 53, 58, and 67 for *Nan-GAL4/+*, *UAS-TrpA1/+,* and *Nan-GAL4 > UAS-TrpA1,* respectively. **(C)** In the *dNf1* background, warming reduced or produced little change in daytime sleep in the genetic controls, whereas activation of Nan+ neurons significantly increased sleep relative to both controls. Despite this increase in sleep quantity, sleep remained highly fragmented (Supplementary Fig. 6 B, C). N = 28, 28, and 62. **(D)** In *dNrx1* mutants, the change in daytime sleep following Nan+ neuron activation was not significantly different from that observed in the corresponding genetic controls. N = 32, 26 and 26. **(E)** In *dFmr1* mutants, activation of Nan+ neurons produced a significant increase in daytime sleep relative to both genetic controls. N = 17, 14, and 21. Points represent individual flies; bars show mean ± SEM. Statistical comparisons were performed using a one-way ANOVA with Holm Šídák’s post hoc comparisons. \*\*\**P* < 0.001; \*\*\*\**P* < 0.0001.

In flies with a control genetic background, increasing the temperature produced a modest increase in sleep in Gal4-only and dTRPA1-only genetic controls (Fig. 4B, left), consistent with known temperature effects (35). In control flies expressing dTRPA1 in Nan+ neurons, however, the temperature shift produced a robust increase in sleep that was significantly greater than the temperature-dependent changes observed in both genetic controls (Fig 4B, right). Thus, consistent with previous studies, direct activation of Nan+ mechanosensory neurons is sufficient to promote sleep in the absence of vibration (9,10).

We first examined dFmr1 mutants, which exhibited abnormal baseline and vibration-evoked CRTC localization in Nan+ neurons (Fig. 3). Thermogenetic activation of Nan+ neurons significantly increased sleep in dFmr1 mutants relative to their genetic controls (Fig. 4 E). Thus, despite altered activity-dependent CRTC signaling in dFmr1 sensory neurons, direct activation of the Nan+ population was sufficient to promote sleep.

In contrast, thermogenetic activation of Nan+ neurons did not significantly increase sleep in dNrx1 mutants relative to their genetic controls (Fig. 4D). This failure occurred despite preserved vibration-evoked CRTC responses in dNrx1 Nan+ neurons (Fig. 3), indicating that neither physiological nor thermogenetic activation of these sensory neurons was sufficient to promote sleep in the dNrx1 background.

dNf1 mutants showed an intermediate response to thermogenetic activation. Increasing the temperature decreased sleep in dNf1 Gal4-only and dTRPA1-only genetic controls, indicating that the effect of warming on sleep was itself altered in the dNf1 background (Fig. 4 C). Nevertheless, activation of Nan+ neurons increased sleep relative to these genetic controls. This increase remained highly fragmented, however, with an increase in sleep bout number but no corresponding increase in bout length (Fig. S6). Thus, direct activation of Nan+ neurons retained the ability to increase sleep in dNf1 mutants but did not restore normal sleep consolidation.

Together, these findings demonstrate that direct activation of Nan+ sensory neurons has different capacities to overcome impaired VIS across NDD models. Nan+ activation promoted sleep in dFmr1 mutants, failed to promote additional sleep in dNrx1 mutants, and increased sleep without restoring normal sleep consolidation in dNf1 mutants. Thus, although all three mutants share a failure of mechanosensory stimulation to promote sleep, direct activation of the sensory neurons underlying this response reveals distinct functional impairments that can converge on the same VIS phenotype.

### Sleep pressure reveals altered state-dependent effects of mechanosensory stimulation in NDD mutants

We next asked whether impaired VIS in NDD mutants was fixed or could be modified by the animal’s homeostatic sleep state. To increase sleep pressure, flies were monitored for 24 h under baseline conditions and then mechanically sleep-deprived for the final 6 h of the night (ZT18– ZT24). Beginning at ZT0, recovery sleep was measured for 3 h under either sham conditions or continuous gentle vibration (Fig. 5 A).

**Figure 5:**
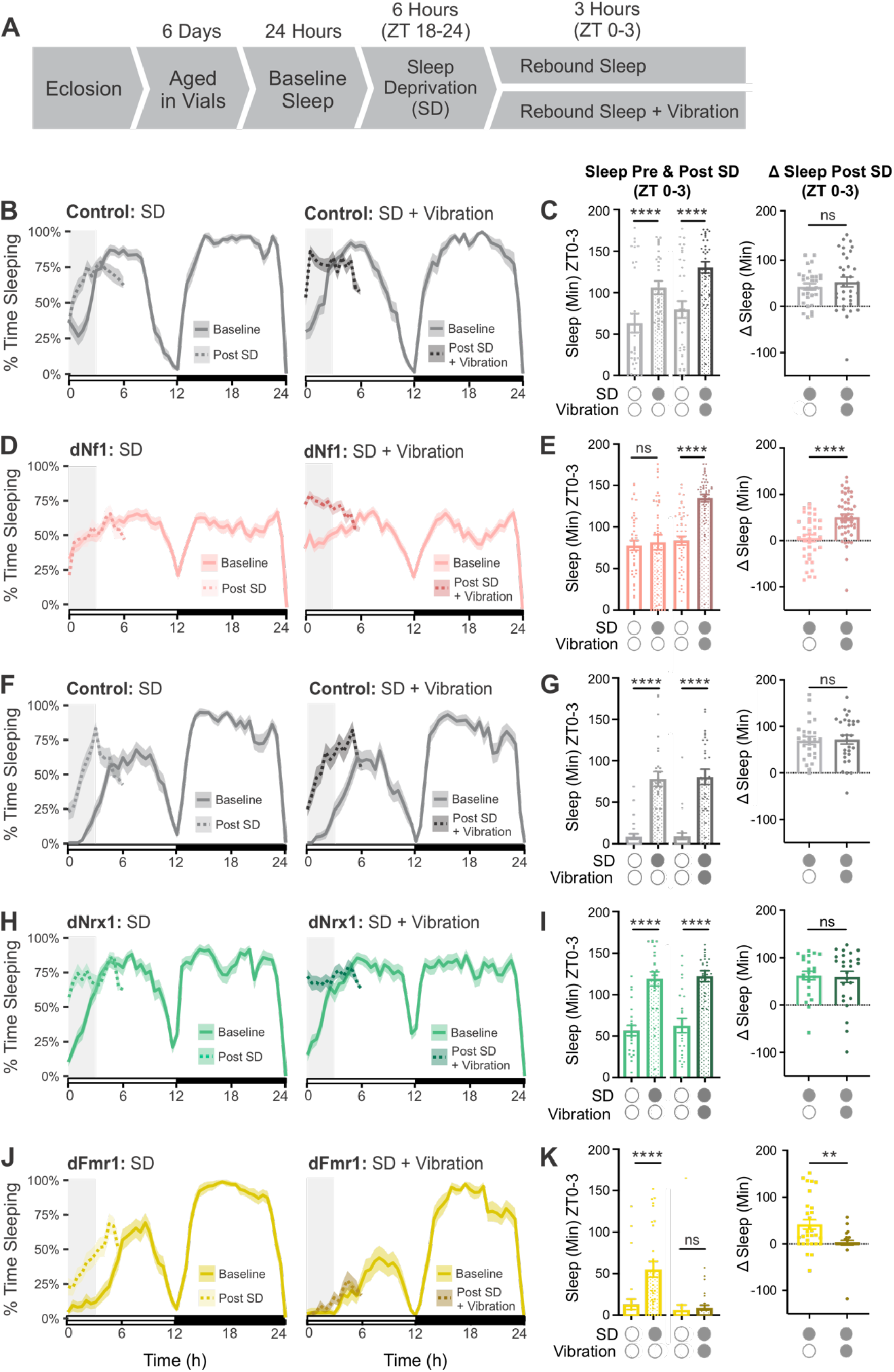
VIS is state-dependent in dNf1 and dFmr1 mutants. **(A)** Experimental design. Following eclosion, female flies were aged for 6 days, and baseline sleep was recorded for 24 h. Flies were then sleep deprived (SD) for 6 h during the latter half of the night (ZT18–24). Rebound sleep was assessed during the first 3 h of the following day (ZT0–3), either under standard conditions or in the presence of gentle vibration. **(B, D, F, H, J)** Averaged sleep traces for the indicated genotypes before and after sleep deprivation. Left panels compare baseline sleep with rebound sleep post-SD; right panels compare baseline sleep with rebound sleep in the presence of vibration. Gray shading denotes the ZT0–3 recovery interval used for quantification. Lines show the mean and shaded regions indicate SEM. **(B)** Genetic control for *dNf1*. SD n = 42, SD + Vibration n = 39 **(D)** *dNf1* mutants. SD n = 38, SD + Vibration n = 49 **(F)** Genetic control for *dNrx1* and *dFmr1*. SD n = 27, SD + Vibration n = 29 **(H)** *dNrx1* mutants. SD n = 23, SD + Vibration n = 25 **(J)** *dFmr1* mutants. SD n = 30, SD + Vibration n = 28. **(C, E, G, I, K)** Quantification of sleep during ZT0–3 for the corresponding genotypes. Left graphs show raw sleep values during baseline and following SD, either with or without vibration during rebound. Right graphs show the change in sleep relative to the matched baseline day (Δ sleep = recovery sleep − baseline sleep). Sleep deprivation increased recovery sleep in both control backgrounds (**C, G**) and in *dNrx1* mutants (**I**), with no significant additional effect of vibration. *dNf1* mutants did not exhibit significant rebound sleep following SD alone, whereas recovery in the presence of vibration significantly increased sleep (**E**). In *dFmr1* mutants, SD increased recovery sleep, but this rebound was abolished when vibration was present (**K**). Points represent individual flies; bars show mean ± SEM. A two-way mixed design repeated measures ANOVA with Holm-Šídák’s multiple comparisons was used to compare raw sleep values and unpaired t-tests were used to compare sleep changes. ns, *P* > 0.05; \**P* < 0.05; \*\**P* < 0.01; \*\*\**P* < 0.001. \*\*\*\**P* < 0.0001

Under sham conditions, sleep-deprived control flies exhibited a significant increase in sleep during the ZT0–ZT3 recovery period relative to the corresponding baseline interval, consistent with a homeostatic rebound following sleep loss (36,37) (Fig. 5 B, C, left). Exposure to vibration did not further augment this response, with similar increases in sleep following deprivation with or without vibration (Fig. 5 B, C, right). Thus, under these conditions, vibration did not add to the homeostatic rebound in control flies.

In *dNf1* mutants, sleep deprivation alone did not produce a significant increase in sleep during the recovery period (Fig. 5 D, E, left). Instead, sleep-deprived *dNf1* flies exhibited fewer sleep bouts and increased waking activity relative to baseline (Supplementary Fig. 7 A, C), indicating an abnormal behavioral response to sleep loss. Strikingly, when sleep-deprived dNf1 mutants were exposed to vibration, sleep increased significantly relative to baseline and to the change observed following sleep deprivation alone (Fig. 5 D, E). This increase was accompanied by a small but significant increase in sleep-bout length (Fig. S7 B). Thus, although neither vibration nor sleep deprivation alone effectively increased sleep in *dNf1* mutants, their combination promoted sleep, demonstrating that the sleep-promoting effect of mechanosensory stimulation can emerge in dNf1 under conditions of elevated homeostatic sleep pressure.

In contrast, *dNrx1* mutants exhibited a significant increase in sleep following sleep deprivation under sham conditions (Fig. 5 H, I, left), indicating a preserved homeostatic rebound. Adding vibration did not further increase recovery sleep (Fig. 5 H, I, right). Thus, unlike dNf1, elevating homeostatic sleep pressure did not reveal a sleep-promoting response to vibration in dNrx1 mutants.

Like *dNrx1* mutants, *dFmr1* mutants exhibited a significant increase in sleep following sleep deprivation under sham conditions, consistent with an intact behavioral rebound response (Fig. 5 J, K, left). Unexpectedly, exposure to vibration nearly eliminated rebound sleep (Fig. 5 J, K, right). Thus, in sleep-deprived *dFmr1* mutants, mechanosensory stimulation opposed rather than augmented the behavioral expression of homeostatic sleep pressure.

Together, these findings demonstrate that the impaired response to mechanosensory stimulation in NDD mutants is strongly influenced by internal sleep state. Elevated sleep pressure revealed a sleep-promoting response to vibration in dNf1 mutants, failed to reveal such a response in dNrx1 mutants, and caused vibration to oppose recovery sleep in dFmr1 mutants. Thus, the shared loss of VIS observed under baseline conditions does not represent a fixed inability of mechanosensory stimulation to influence sleep. Rather, these mutations differentially alter how mechanosensory input interacts with homeostatic sleep drive to shape behavioral state.

## DISCUSSION

Sleep disturbances are common in neurodevelopmental disorders (NDDs), yet their expression is notably heterogeneous, both across disorders and among individuals with the same genetic diagnosis (16–18,38). This heterogeneity is typically described in terms of conventional sleep phenotypes, including differences in sleep duration, timing, and fragmentation. Here, we find three *Drosophila* models of NDDs with markedly different baseline sleep phenotypes converging on a shared impairment in the ability of prolonged mechanosensory stimulation to promote sleep. These findings suggest that examining how sleep responds to sensory context in NDDs may reveal shared dysfunction that is not apparent from measures of baseline sleep alone.

The convergence on impaired VIS was particularly notable because it did not arise from a common underlying defect. Across behavioral, cellular, and thermogenetic assays, the three NDD models showed evidence for disruption at different points along the pathway linking mechanosensory input to sleep. Thus, distinct perturbations of sensory processing or its ability to influence sleep can produce the same systems-level phenotype. This mechanistic diversity suggests that sensory-sleep integration may represent a point of phenotypic convergence across molecularly distinct forms of neurodevelopmental dysfunction.

Our findings further indicate that sensory-sleep dysfunction is not a fixed phenotype, but depends on the animal’s internal state. Under elevated homeostatic sleep pressure, vibration promoted sleep in dNf1 mutants, continued to fail to promote sleep in dNrx1 mutants, and opposed homeostatic recovery sleep in dFmr1 mutants. Thus, the expression of sensory-sleep dysfunction reflects an interaction among genotype, sensory context, and internal sleep state. This state dependence provides another potential source of phenotypic variability that may be missed when sleep is assessed only under baseline conditions.

The combination of behavioral, cellular, and functional manipulations in this study provides insight into how sensory-sleep integration is disrupted in each NDD model. Although all three mutations converged on impaired VIS, their responses to vibration, activity-dependent CRTC signaling in Nan+ sensory neurons, direct Nan+ activation, and elevated sleep pressure revealed distinct points of dysfunction along the sensory-sleep axis. Together, these findings reveal distinct routes through which each mutation can disrupt the transformation of sensory input into sleep.

### *dNf1*: mechanosensory input promotes sleep in a state-dependent manner

Among the three models, the *dNf1* phenotype most strongly implicates a defect downstream of initial sensory processing. *dNf1* mutants exhibited a normal acute behavioral response to vibration (Fig. 2 B) and a normal vibration-associated increase in CRTC nuclear localization in Nan+ sensory neurons (Fig. 3 B). These findings indicate that vibration is detected behaviorally and engages activity-associated signaling in the Nan+ population, making deficient primary sensory-neuron activation unlikely to fully explain the loss of VIS. The preservation of mechanosensory responses in *dNf1* contrasts with the effects of *Nf1* loss in other sensory contexts. *Nf1* loss can attenuate pheromone-evoked activity in Ppk23+ chemosensory neurons (39), while *Nf1* mutant larvae exhibit behavioral hypersensitivity to tactile stimulation and altered neurotransmission at the neuromuscular junction (40). Together, these findings suggest that the consequences of *Nf1* loss on sensory processing depend on sensory modality, neuronal population, and developmental stage, consistent with the heterogeneous sensory-processing differences reported in children with NF1 (25).

Thermogenetic activation provided further evidence that the capacity of Nan+ sensory neurons to promote sleep remains at least partially intact in dNf1 mutants. Direct Nan+ neuron activation increased total sleep relative to the corresponding dNf1 genetic controls (Fig. 4 C). However, the underlying fragmented sleep architecture persisted, with an increase in sleep-bout number but no increase in bout length (Fig. S6), consistent with the baseline sleep phenotype of dNf1 mutants (26,27). Thus, direct activation of Nan+ neurons can promote additional sleep in dNf1 mutants but does not overcome the broader sleep-fragmentation phenotype associated with loss of Nf1.

The response to sleep deprivation provides an additional perspective on sleep regulation in *dNf1* mutants. In the absence of vibration, sleep-deprived *dNf1* mutants did not display a conventional increase in rebound sleep (Fig. 5 D, E), instead showing fewer sleep bouts and increased waking activity (Fig. S7). This abnormal response could reflect impaired accumulation of homeostatic sleep pressure or an inability to translate increased sleep need into increased sleep. Previous work using pan-neuronal *Nf1* knockdown found that sleep deprivation does produce rebound sleep, although this rebound remains highly fragmented (41), suggesting that homeostatic sleep drive can persist in the context of *Nf1* dysfunction. Strikingly, sleep-deprived *dNf1* mutants in our experiments slept more when recovery occurred in the presence of vibration (Fig. 5 D, E). Thus, although neither sleep deprivation nor vibration alone increased sleep under the conditions tested, combining elevated sleep pressure with mechanosensory stimulation was sufficient to do so.

Together, these findings suggest that the loss of VIS in *dNf1* mutants does not reflect a simple failure to detect or respond to mechanosensory input. Nan+ sensory neurons respond to vibration, direct activation of these neurons can increase sleep, and vibration can promote sleep when homeostatic sleep pressure is elevated. The latter finding is particularly notable because neither vibration nor sleep deprivation alone increased sleep under the conditions tested, suggesting that sensory and homeostatic influences can converge to promote sleep in *dNf1* mutants. How *Nf1* loss disrupts this interaction remains unclear, but these findings indicate that the failure to express VIS under baseline conditions does not represent a complete loss of the capacity for mechanosensory input to promote sleep.

### *dNrx1*: impaired coupling of mechanosensory activity to sleep

With loss of *dNrx1*, our results implicate a disruption between mechanosensory processing and its behavioral consequences. At the vibration intensity used for VIS, *dNrx1* mutants exhibited reduced acute behavioral responsiveness to vibration (Fig. 2 D, F). At higher vibration intensities, however, behavioral response rates were comparable to controls (Fig. S4 B), demonstrating that *dNrx1* mutants remain capable of detecting and behaviorally responding to vibration under these conditions. Thus, the impaired VIS phenotype is unlikely to reflect a complete loss of mechanosensory responsiveness or an inability to generate a locomotor response.

Despite their reduced acute behavioral responsiveness, Nan+ neurons in *dNrx1* mutants exhibited a significant vibration-associated increase in nuclear CRTC (Fig. 3 B). Although CRTC localization is an indirect activity-associated readout and does not measure action potentials or neurotransmitter release directly, this response indicates that vibration engages activity-associated signaling within the Nan+ sensory population. The dissociation between preserved vibration-evoked CRTC signaling and reduced behavioral responsiveness suggests that the *dNrx1* phenotype cannot be explained by a failure to activate Nan+ sensory neurons.

This interpretation is further supported by the thermogenetic experiments. Direct activation of Nan+ neurons did not increase sleep in *dNrx1* mutants compared to controls (Fig. 4 D), indicating that increasing Nan+ neuron excitability is insufficient to promote sleep in this background. Elevating homeostatic sleep pressure similarly failed to reveal a sleep-promoting response to vibration (Fig. 5). Importantly, however, *dNrx1* mutants exhibited rebound sleep following deprivation, demonstrating that they retain the capacity to increase sleep in response to homeostatic drive. Together, these findings suggest that the *dNrx1* defect lies not in the general capacity to generate additional sleep, but in the ability of mechanosensory input, including direct activation of Nan+ neurons, to promote sleep.

A defect in synaptic transmission or downstream circuit connectivity would be consistent with the known functions of Neurexin. Neurexins are presynaptic adhesion molecules that organize trans-synaptic signaling and regulate synapse-specific properties; in mammals, loss of neurexin function disrupts presynaptic Ca²⁺-channel organization, Ca²⁺-triggered vesicle release, and circuit-specific excitatory and inhibitory connectivity (42–45). In *Drosophila*, Neurexin is required for normal active-zone organization, synaptic-vesicle distribution, and evoked neurotransmission (46,47). Loss of *dNrx1* causes ectopic accumulation of synaptic-vesicle and active-zone proteins along axons and disrupts synaptic organization and function (46). These established roles suggest that the dissociation we observe between Nan+ sensory-neuron activation and sleep could arise from impaired transmission of sensory information to downstream targets. In this model, Nan+ neurons remain responsive to vibration, but their activity is not effectively translated into the downstream signals required to promote sleep. Future work will aim to determine whether *dNrx1* alters the synaptic output or connectivity of Nan+ neurons.

### *dFmr1*: abnormal sensory encoding and state-dependent inversion of the vibration response

The *dFmr1* phenotype differs from both *dNf1* and *dNrx1* because abnormal activity-associated signaling was evident within Nan+ sensory neurons. Under baseline conditions, Nan+ neurons in *dFmr1* mutants exhibited elevated nuclear CRTC relative to controls, and vibration produced a decrease in nuclear CRTC rather than the increase observed in control flies (Fig. 3 B). Behaviorally, *dFmr1* mutants also exhibited reduced acute responsiveness at the standard vibration intensity (Fig. 2 E, F), although behavioral responses were observed at higher stimulus intensities (Fig. S4 C). Thus, the impaired response to vibration does not reflect an absolute inability to detect mechanosensory stimulation, but is accompanied by abnormal basal and vibration-evoked CRTC signaling within Nan+ sensory neurons.

The elevated basal CRTC signal and its decrease following vibration indicate that activity-associated signaling in Nan+ neurons is altered in *dFmr1* mutants, although the physiological basis of these changes remains unclear. CRTC localization is regulated by activity-dependent signaling pathways, including calcineurin, cAMP, and SIK-dependent signaling (34,48), and therefore does not provide a direct measure of membrane excitability or firing rate. The abnormal CRTC response could reflect altered neuronal activity, adaptation to sensory stimulation, or changes in the intracellular pathways regulating CRTC itself. Notably, FMRP has been implicated in regulation of calcium homeostasis and stimulus-evoked cAMP signaling across fly, mouse, and human Fragile X models (49–52), providing several potential mechanisms through which loss of *Fmr1* could alter activity-dependent signaling in these sensory neurons.

In contrast to *dNrx1*, direct thermogenetic activation of Nan+ neurons significantly increased sleep in *dFmr1* mutants (Fig. 4 E), demonstrating that increasing activity in this sensory population retains the capacity to promote sleep. Sleep deprivation also produced a conventional rebound under sham conditions (Fig. 5 J, K), indicating that *dFmr1* mutants remain capable of increasing sleep in response to homeostatic drive. Together, these findings argue against a general inability to generate additional sleep drive and distinguish the *dFmr1* phenotype from the failure of direct Nan+ activation to promote sleep in *dNrx1*. Instead, the impaired VIS phenotype in *dFmr1* occurs despite preserved sleep-promoting responses to both direct Nan+ activation and elevated homeostatic sleep pressure.

The effect of vibration following sleep deprivation revealed an additional state-dependent abnormality. Although *dFmr1* mutants exhibited rebound sleep under sham conditions, vibration nearly eliminated this response (Fig. 5 J, K). Thus, under elevated homeostatic sleep pressure, mechanosensory stimulation did not simply fail to promote sleep but instead opposed the expression of recovery sleep. This finding further distinguishes the *dFmr1* phenotype from a general defect in homeostatic sleep regulation and demonstrates that the behavioral consequence of mechanosensory stimulation depends strongly on internal sleep state. Our results suggest that loss of *Fmr1* alters how sensory input interacts with homeostatic sleep drive.

Together, these findings suggest that the loss of VIS in *dFmr1* mutants is associated with an abnormal response to mechanosensory input rather than a general inability to increase sleep. Nan+ sensory neurons exhibit abnormal basal and vibration-evoked CRTC signaling, yet direct activation of these neurons remains capable of promoting sleep, and homeostatic sleep pressure produces a normal rebound in the absence of vibration. Moreover, the effect of vibration itself depends on internal state, failing to promote normal VIS under baseline conditions and opposing recovery sleep following sleep deprivation. Thus, loss of *dFmr1* disrupts both the cellular response to mechanosensory stimulation and the state-dependent relationship between sensory input and sleep.

### Limitations

Several limitations should be considered. Nan-GAL4 labels a broad population of Nan-expressing sensory neurons, including chordotonal neurons in the antennae and legs, and does not selectively isolate the functionally distinct vibration-sensitive subgroups of Johnston’s organ (31,32,53). CRTC nuclear localization provides an indirect measure of recent activity-associated signaling rather than a direct measure of neuronal firing, and genotype-dependent changes in the signaling pathways regulating CRTC could alter its localization independently of neuronal activity (34,48). Similarly, thermogenetic activation does not reproduce the spatial or temporal patterns of activity evoked by natural vibration, and the temperature shift itself affected sleep differently across genotypes. Our experiments also do not directly measure transmission from Nan+ sensory neurons to downstream sleep-regulatory populations, limiting our ability to anatomically localize the functional defects identified in each mutant. Finally, the mutations examined here were present throughout the animal, preventing assignment of these phenotypes to cell-autonomous functions within sensory or sleep-regulatory neurons. These considerations limit precise anatomical and molecular localization of the underlying defects but do not alter the central finding that distinct neurodevelopmental mutations converge on impaired sensory-sleep integration.

### Conclusions

Our findings suggest that shared sleep dysfunction across NDDs may be most apparent in how sleep responds to sensory context rather than in baseline measures of sleep itself. The three NDD models studied here exhibit markedly different baseline sleep phenotypes, yet converge on impaired sensory-sleep integration, with distinct disruptions along the sensory-sleep axis producing the same behavioral endpoint. Moreover, the expression of this shared phenotype depends on internal sleep state, emphasizing that behavioral phenotypes emerge from interactions among genotype, internal state, and environmental context. Thus, heterogeneity in conventional measures such as sleep duration or fragmentation may obscure shared dysfunction in the processes that dynamically regulate sleep. Identifying such systems-level phenotypes may provide a path toward understanding biological convergence across genetically and phenotypically diverse neurodevelopmental disorders.

## METHODS

### Drosophila husbandry

Flies were raised and maintained on standard molasses-based food containing 8.0% molasses, 0.55% agar, 0.2% Tegosept, and 0.5% propionic acid. Unless otherwise indicated, flies were maintained at 25°C under a 12-h light:12-h dark (LD) cycle. For thermogenetic experiments, flies were raised at 22°C to minimize basal activation of the thermosensitive effector.

Nan-GAL4 (BDSC #24903) was obtained from the Bloomington Drosophila Stock Center. dNf1 mutants were trans-heterozygous for the *Nf1^P1^* and *Nf1^P2^* mutant alleles (54), which were generously provided by Dr. Amita Sehgal. The parental P-element line K33 served as the genetic control for experiments using the original *Nf1* mutant stocks. dNrx1 mutants were trans-heterozygous for the *Nrx1^273^* and *Nrx1^241^* mutant alleles (46), which were generously provided by Dr. Thomas Jongens. *dFmr1* mutants were homozygous for the null allele *dFmr1^3^* (55) and were also provided by Dr. Thomas Jongens. The w^1118^ (iso31Bw−) strain served as the genetic-background control for the dNrx1 and dFmr1 mutant stocks.

For experiments requiring newly generated stocks carrying *Nan-GAL4*, *UAS-CRTC::GFP*, or *UAS-dTRPA1*, the experimental and control lines were constructed in a common isogenic *(iso31)* genetic background. Transgene-matched flies carrying the corresponding GAL4 driver and/or UAS effector but lacking the mutant alleles were used as controls, allowing the same background-matched control genotype to be compared across mutant groups where appropriate.

### Sleep assays

Adult female flies were collected 1–2 d after eclosion and maintained in groups until 4–7 d of age at 25°C under a 12-h:12-h LD cycle, unless otherwise indicated. Females were housed with males before loading to ensure that they were mated. Flies were briefly anesthetized with CO₂ using fly pads (Genesee Scientific, catalog no. 59-114) and individually loaded into glass activity tubes containing food composed of 5% sucrose and 2% agar. Tubes were placed in multibeam Drosophila Activity Monitoring systems (DAM5H; Trikinetics), and flies were acclimated in light-and temperature-controlled incubators for at least 24 h before data collection. When male flies were used (Fig. S3), male flies were collected, housed, and loaded in a similar manner as females.

Experiments were conducted at 25°C, except for thermogenetic experiments. For thermogenetic activation, flies were monitored for 24 h at 22°C to establish baseline sleep and subsequently for 24 h at 29°C to activate dTRPA1-expressing neurons. Locomotor activity was recorded in 1-min bins. Sleep was defined as a period of at least 5 consecutive min without a detected beam crossing. Waking activity was calculated as the number of beam crossings per minute of wakefulness. Data were analyzed using the Rethomics framework, and daytime and nighttime sleep were quantified separately in 12-h bins.

For vibration-induced sleep experiments, baseline sleep was recorded for 24 h. Beginning at zeitgeber time 0 (ZT0), flies were then exposed to continuous gentle vibration for 24 h, as previously described (9). Briefly, activity monitors were positioned on a shelf approximately 40 cm above an analog multi-tube vortexer (Fisher Scientific, Cat#: 02-215-450) set to an intensity of 3. The onset and duration of mechanical stimulation were controlled using an LC4 Light Controller (Trikinetics).

For sleep-deprivation experiments, baseline sleep was recorded for 24 h in multibeam activity monitors. On the following day, the glass tubes containing individual flies were transferred to single-beam activity monitors (DAM2; Trikinetics) and securely mounted on a mechanical vortexer (Fisher Scientific, Cat#: 02-215-450). Flies were sleep-deprived from ZT18 to ZT24 by administering a 2-s mechanical stimulus at a randomly selected time within a 1-min interval. At ZT0, the activity tubes were returned to multibeam monitors and assigned to vibration or sham conditions. Flies in the vibration condition were positioned on a shelf approximately 40 cm above the vortexer and exposed to continuous gentle vibration for 6 h, as described above. Sham-treated flies underwent the same sleep-deprivation and handling procedures but were transferred to a separate incubator and were not exposed to continuous vibration. Sleep during the first 3 h of recovery (ZT0-3) was used for analysis.

### Antennectomy

Antennae were removed as previously described (9). Briefly, 3-day-old female flies were anesthetized with CO₂, and all three antennal segments were removed using fine forceps (Fine Science Tools, Foster City, CA). Following surgery, flies were returned to food vials and allowed to recover for 3 days before sleep experiments. Intact controls underwent the same anesthesia and handling procedures without antennal removal.

### Locomotor responses to vibration

Adult female flies were collected and loaded into multibeam activity monitors as described above. Following acclimation, baseline locomotor activity was recorded for 30 min beginning at ZT3. Flies were subsequently exposed to 30 min of gentle vibration using the analog multi-tube vortexer at an intensity setting of 3, as described above. For vibration at varying intensities, flies were exposed to 20Hz vibration using a Universal Vibration Stage (UVS-01; Tau Scientific) with settings of 0.18 for low intensity assays and 0.31 for high intensity assays. To minimize the influence of sleep state on the response to vibration, flies that were asleep at vibration onset were excluded from the analysis.

Activity was recorded using the “counts” acquisition mode, which captures both local movements occurring within an individual beam and movements between adjacent beams. Activity counts were initially collected in 1-min bins and averaged into 5-min bins for analysis and visualization. The 15 min immediately preceding vibration onset were used to define baseline activity. A fly was classified as a vibration “responder” if its absolute change in activity during the first 5 min of vibration exceeded two standard deviations of its baseline activity.

### CRTC localization in Nan+ antennal sensory neurons

To visualize CRTC in Nan+ sensory neuron cell bodies, mutant lines carrying *dNf1*, *dNrx1*, or *dFmr1* mutations and the *Nan-Gal4* driver were crossed to the corresponding mutant lines carrying *UAS-CRTC-GFP* and *UAS-mCD8::mCherry*. This strategy enabled visualization of CRTC-GFP together with mCD8::mCherry-labeled neuronal membranes in each mutant background.

Adult flies were briefly anesthetized with CO₂, individually loaded into multibeam activity monitors (DAM5H system; Trikinetics), and acclimated for 24 h. Flies were maintained at 25°C under a 12-h light:12-h dark cycle. At ZT3, the monitors were placed on a Universal Vibration Stage (UVS-01; Tau Scientific). Flies in the vibration condition were exposed to 20-Hz vibration at an instrument intensity setting of 0.24 for 5 min, whereas sham-treated flies were placed on the stage without vibration.

Immediately afterwards, flies were individually removed from the activity tubes, briefly immersed in 200-proof ethanol, and transferred to 4% paraformaldehyde for 1 h. Fixed flies were washed three times for 5 min each in phosphate-buffered saline (PBS). One antenna from each fly was then dissected in PBS using fine forceps. Dissected antennae were dehydrated by three 5-min washes in methanol and subsequently rehydrated through a graded methanol series in PBS containing 0.5% Triton X-100 (PBST; 75%, 50%, and 25% methanol). Samples were incubated for 48 h at 4°C in primary antibody solution containing chicken anti-GFP (1:400; Invitrogen) and mouse anti-mCherry (1:200; Developmental Studies Hybridoma Bank) diluted in 0.5% PBST. Antennae were washed four times for 15 min each in 0.5% PBST and incubated for 48 h at 4°C with goat anti-chicken Alexa Fluor 488 (1:250; Invitrogen) and donkey anti-mouse Alexa Fluor 594 (1:250; Invitrogen) diluted in 0.5% PBST.

Following secondary antibody incubation, samples were washed twice for 15 min in PBST and twice for 15 min in PBS. Antennae were then equilibrated through a graded glycerol series consisting of 40%, 60%, and 80% glycerol in PBS for 10 min at each concentration. Samples were mounted on glass slides in VECTASHIELD mounting medium, covered with glass coverslips, and imaged immediately using a Leica TCS SP8 confocal microscope. The second antennal segment was imaged using a 63× oil-immersion objective with a z-step size of 0.5 µm. Identical acquisition settings were used across experimental groups.

Images were processed and analyzed in Fiji/ImageJ (National Institutes of Health) by an experimenter blinded to genotype and treatment condition. Five Nan+ neurons were randomly selected from each antenna, and separate regions of interest (ROIs) were drawn around the nucleus and cytoplasm of each neuron. The subcellular distribution of CRTC-GFP was quantified by calculating a nuclear localization index (NLI) for each neuron according to the following formula: NLI = (mean nuclear GFP intensity - mean cytoplasmic GFP intensity)/(mean nuclear GFP intensity + mean cytoplasmic GFP intensity) (34). The NLI values from the five neurons were averaged to obtain a single mean NLI for each antenna, which was treated as the unit of analysis.

### Statistical Analysis

Statistical analyses were performed using GraphPad Prism version 11 (GraphPad Software, Boston, MA, USA). Unless otherwise stated, each fly represented an independent biological replicate. For CRTC imaging experiments, measurements from five neurons within one antenna were averaged to produce one value per fly. Individual data points are shown where possible, and summary data are presented as mean ± SEM. All statistical tests were two-sided, and *P* < 0.05 was considered statistically significant. Flies that died during an experiment were excluded from analysis. Potential outliers in raw-value datasets were identified using the robust regression and outlier removal (ROUT) method with Q = 0.1%.

Normality was assessed using the Shapiro–Wilk test with α = 0.05. Comparisons between two independent groups were performed using unpaired *t*-tests for normally distributed data or Mann-Whitney tests when the normality assumption was not met. These tests were used to compare baseline sleep parameters and vibration-induced changes in sleep between mutants and their corresponding genetic controls. To determine whether vibration significantly altered sleep within each genotype, vibration-induced sleep values were also compared with a hypothetical mean of zero using one-sample *t*-tests.

To determine whether differences in baseline sleep accounted for genotype-dependent changes in VIS, vibration-day daytime sleep was analyzed separately for each mutant–control comparison using ordinary least-squares multiple linear regression. Baseline daytime sleep was entered as a continuous predictor, genotype as a categorical predictor, and vibration-day daytime sleep as the outcome. Initial models included a baseline sleep-by-genotype interaction to test whether the relationship between baseline and vibration-day sleep differed by genotype. Because the interaction was not significant in any comparison, interaction terms were removed and reduced models containing baseline sleep and genotype as main effects were analyzed. Simple linear regression was used to visualize the relationship between baseline and vibration-day sleep and to estimate the pooled slope when genotype-specific slopes did not differ significantly.

For antennectomy experiments, baseline daytime sleep and vibration-induced sleep were analyzed separately for each mutant–control comparison using ordinary two-way ANOVA, with genotype (mutant versus control) and antennectomy status (intact versus antennectomized) as between-subject factors. The genotype-by-antennectomy interaction was evaluated for each outcome. When the interaction was significant, uncorrected Fisher’s least significant difference tests were used for planned pairwise comparisons.

Vibration-evoked locomotor activity was evaluated with paired t-tests. When the assumptions of the parametric analysis were not met, Wilcoxon matched-pairs signed rank test was used. Differences in the proportion of flies classified as responders to vibration onset were evaluated using two-sided Fisher’s exact tests. Flies that were asleep at vibration onset were excluded from the locomotor-response analysis.

CRTC nuclear localization indices were analyzed using an ordinary two-way ANOVA, with genotype and vibration condition (sham or vibration) as between-subject factors. The genotype-by-condition interaction was evaluated, followed by Holm-Šídák multiple-comparisons tests comparing sham and vibration conditions within each genotype and comparing genotypes within each condition.

For thermogenetic experiments, the change in daytime sleep produced by the temperature shift was calculated for each fly as daytime sleep at 29°C minus daytime sleep at 22°C. Within each genetic background, changes among the GAL4-only, UAS-TrpA1-only, and *Nan-GAL4 > UAS-TrpA1* groups were compared using ordinary one-way ANOVA followed by Holm-Šídák multiple-comparisons tests. Raw values for daytime sleep, sleep-bout number, mean sleep-bout length, and waking activity index were analyzed separately within each genetic background using two-way mixed-design ANOVA. Transgene genotype was treated as a between-subject factor, and temperature (22°C versus 29°C) was treated as a within-subject factor. When the transgene genotype-by-temperature interaction was significant, Holm-Šídák multiple-comparisons tests were used to compare measurements at 22°C and 29°C within each transgene genotype.

For sleep-deprivation experiments, raw sleep during the ZT0-ZT3 recovery interval was analyzed separately for each genotype using a two-way mixed-design repeated-measures ANOVA. Recording condition (baseline versus post-sleep deprivation) was treated as a within-subject factor, and recovery treatment (sham versus vibration) was treated as a between-subject factor. Holm–Šídák multiple-comparisons tests were used for planned comparisons. The change in sleep was calculated for each fly as recovery sleep minus sleep during the corresponding baseline interval. Changes in sleep between the sham and vibration recovery groups were compared using unpaired t-tests or Mann-Whitney tests, as appropriate.

