## Supplemental Figures for "Sensory-sleep dysfunction is a shared phenotype across genetically distinct neurodevelopmental disorder models"

### SUPPLEMENTARY MATERIALS

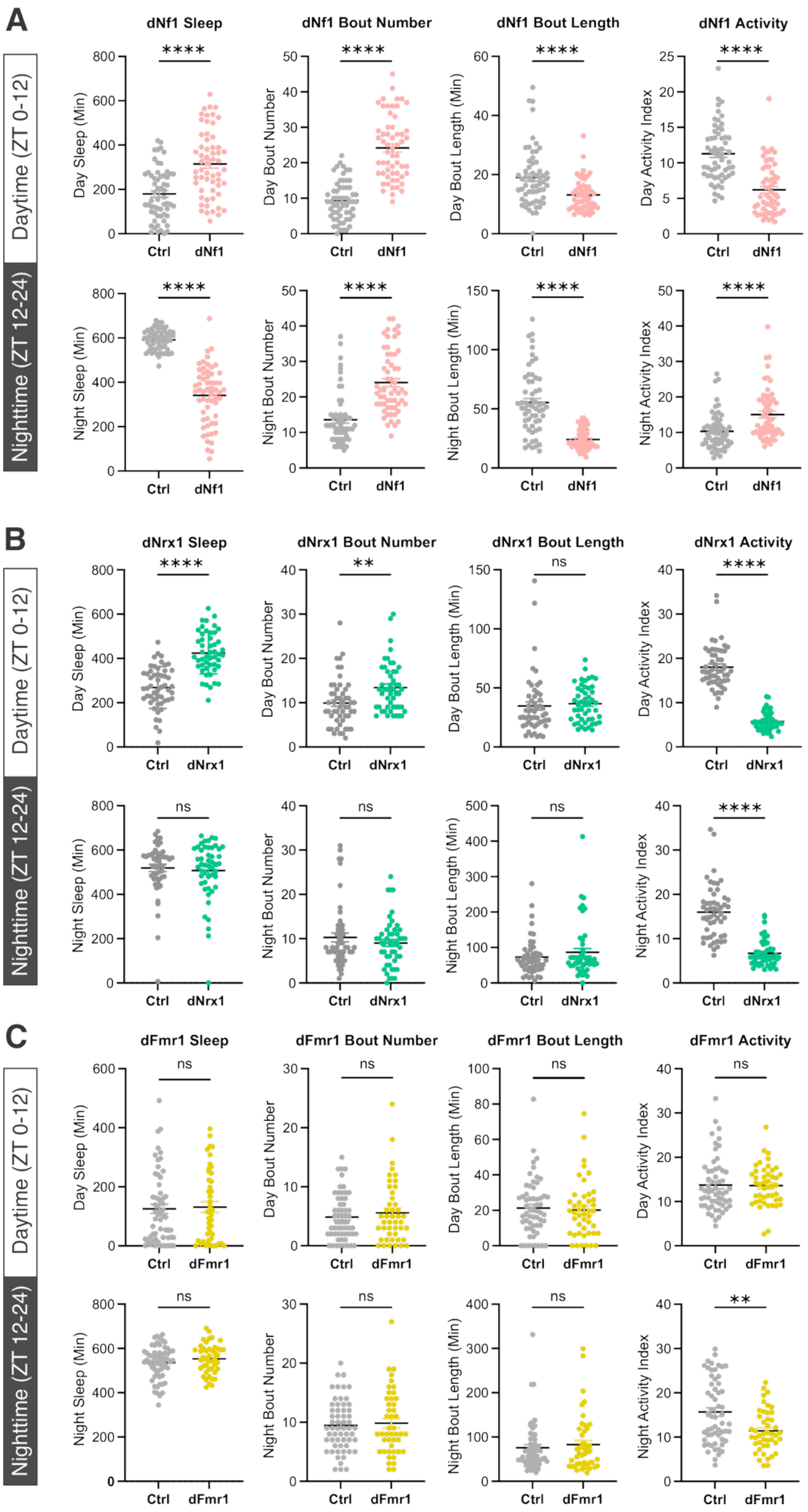
A
Daytime (ZT 0-12)
Nighttime (ZT 12-24)
dNf1 Sleep
dNf1 Bout Number
dNf1 Bout Length
dNf1 Activity
Day Sleep (Min)
Day Bout Number
Day Bout Length (Min)
Day Activity Index
Night Sleep (Min)
Night Bout Number
Night Bout Length (Min)
Night Activity Index
****
Ctrl
dNf1
B
dNrx1 Sleep
dNrx1 Bout Number
dNrx1 Bout Length
dNrx1 Activity
****
**
ns
dNrx1
C
dFmr1 Sleep
dFmr1 Bout Number
dFmr1 Bout Length
dFmr1 Activity
dFmr1

**Supplementary Figure 1. Baseline sleep and waking activity phenotypes differ across NDD mutant models.**

Baseline daytime (ZT0–12; upper rows) and nighttime (ZT12–24; lower rows) sleep parameters were quantified for each mutant and its corresponding genetic control. From left to right, graphs show total sleep, sleep-bout number, mean sleep-bout length, and waking activity index. Each point represents an individual fly; bars show mean ± SEM. **(A)** Baseline sleep in *dNf1* mutants and their genetic controls. During both the day and night, *dNf1* mutants exhibited increased sleep-bout number and reduced bout length, indicating severe sleep fragmentation. Compared with controls, *dNf1* mutants slept more and exhibited reduced waking activity during the day, whereas they slept less and exhibited increased waking activity during the night. Control, *n* = 59; *dNf1*, *n* = 58. **(B)** Baseline sleep in *dNrx1* mutants and their genetic controls. *dNrx1* mutants exhibited increased daytime sleep and daytime bout number, without a significant difference in daytime bout length. Nighttime sleep, bout number, and bout length did not differ significantly between genotypes. Waking activity was reduced in *dNrx1* mutants during both the day and night. Control, *n* = 52; *dNrx1*, *n* = 49. **(C)** Baseline sleep in *dFmr1* mutants and their genetic controls. Daytime and nighttime sleep amount, bout number, and bout length did not differ significantly between *dFmr1* mutants and controls. Daytime waking activity was also comparable between genotypes, whereas nighttime waking activity was modestly reduced in *dFmr1* mutants. Control, *n* = 58; *dFmr1*, *n* = 45. Control and mutant groups were compared using unpaired t-tests. ns, $P > 0.05$; $^{**}P < 0.01$; $^{****}P < 0.0001$.

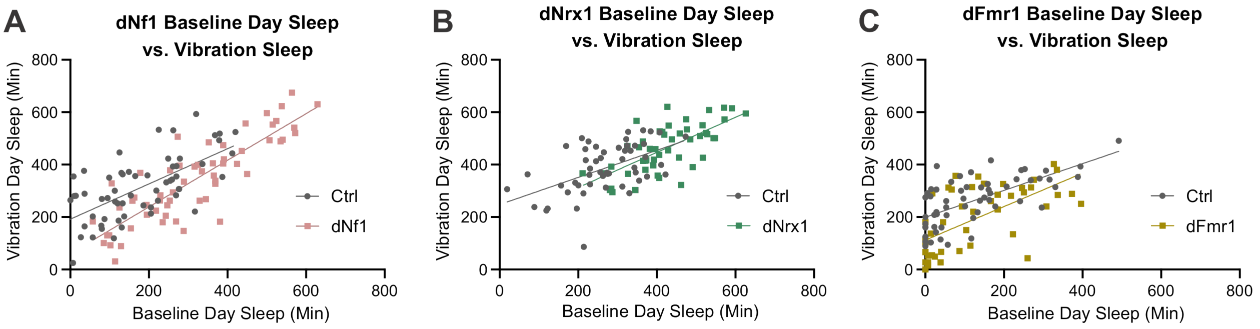


**Supplementary Figure 2. Baseline sleep differentially contributes to vibration-day sleep across NDD models.**

The relationship between baseline daytime sleep and daytime sleep during vibration was examined for *dNf1* (**A**), *dNrx1* (**B**), and *dFmr1* (**C**) mutants and their corresponding genetic controls. Each point represents an individual fly, and lines show the best-fit linear regressions for each genotype. Vibration-day sleep was analyzed separately for each mutant–control comparison using multiple linear regression, with baseline sleep, genotype, and the baseline-by-genotype interaction as predictors. The interaction was not significant for *dNf1* ($P = 0.0833$), *dNrx1* ($P = 0.3436$), or *dFmr1* ($P = 0.3523$); reduced models containing baseline sleep and genotype as main effects were therefore analyzed. Baseline sleep significantly predicted vibration-day sleep in all three comparisons (all $P = 0.0001$). Genotype remained a significant predictor in the *dNf1* and *dFmr1* comparisons (both $P = 0.0001$), with pooled slopes of 0.8075 and 0.5671, respectively. In contrast, genotype was not a significant predictor in the *dNrx1* comparison ($P = 0.4133$; pooled slope = 0.5905). **(A)** Control, *n* = 59; *dNf1*, *n* = 58. **(B)** Control, *n* = 52; *dNrx1*, *n* = 49. **(C)** Control, *n* = 58; *dFmr1*, *n* = 45.

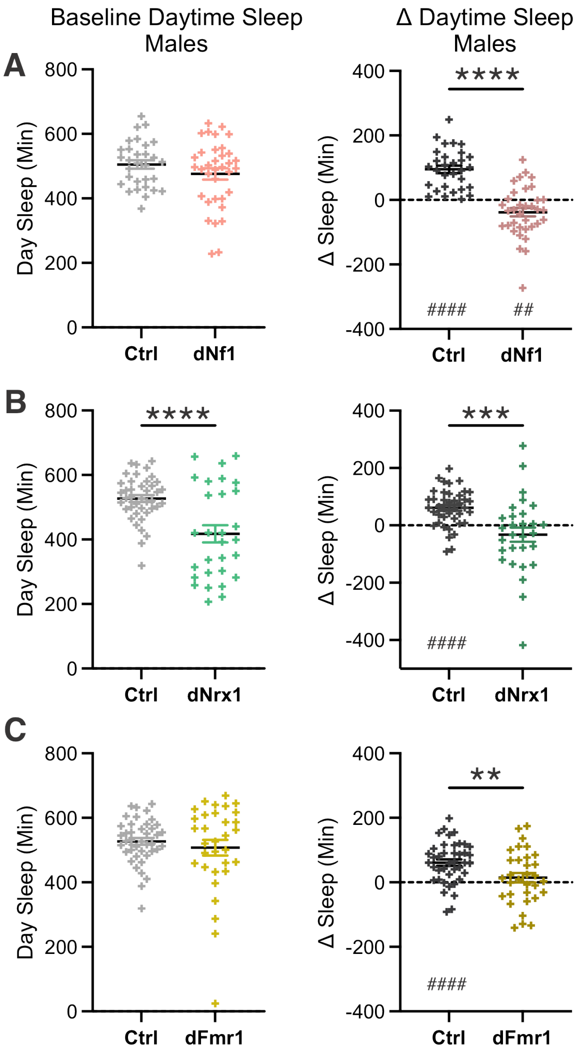


**Supplementary Figure 3. Vibration-induced sleep is impaired in male NDD mutants.** Baseline daytime sleep (left) and vibration-induced sleep (right) were quantified in male *dNf1* (**A**), *dNrx1* (**B**), and *dFmr1* (**C**) mutants and their corresponding genetic controls. Baseline daytime sleep was significantly lower in dNrx1 mutant males than controls. VIS was calculated for each fly as the change in daytime sleep between the vibration and baseline days (Δ sleep = vibration-day sleep − baseline-day sleep, ZT0–12). Control males exhibited significant increases in daytime sleep during vibration. In contrast, *dNrx1* and *dFmr1* males showed no significant change in daytime sleep, whereas *dNf1* males exhibited a significant reduction in sleep. The change in daytime sleep was significantly smaller in all three mutants than in their respective controls. Each point represents an individual fly; bars show mean ± SEM. **(A)** Control, $n = 30$; *dNf1*, $n = 35$. **(B)** Control, $n = 41$; *dNrx1*, $n = 30$. **(C)** Control, $n = 41$; *dFmr1*, $n = 32$. Mutants and controls were compared using unpaired *t*-tests, indicated by asterisks. Changes within each genotype were compared with a hypothetical mean of zero using one-sample *t*-tests, indicated by number signs. $^{**}P < 0.01$; $^{***}P < 0.001$; $^{****}P < 0.0001$; $^{\#\#}P < 0.01$; $^{\#\#\#\#}P < 0.0001$.

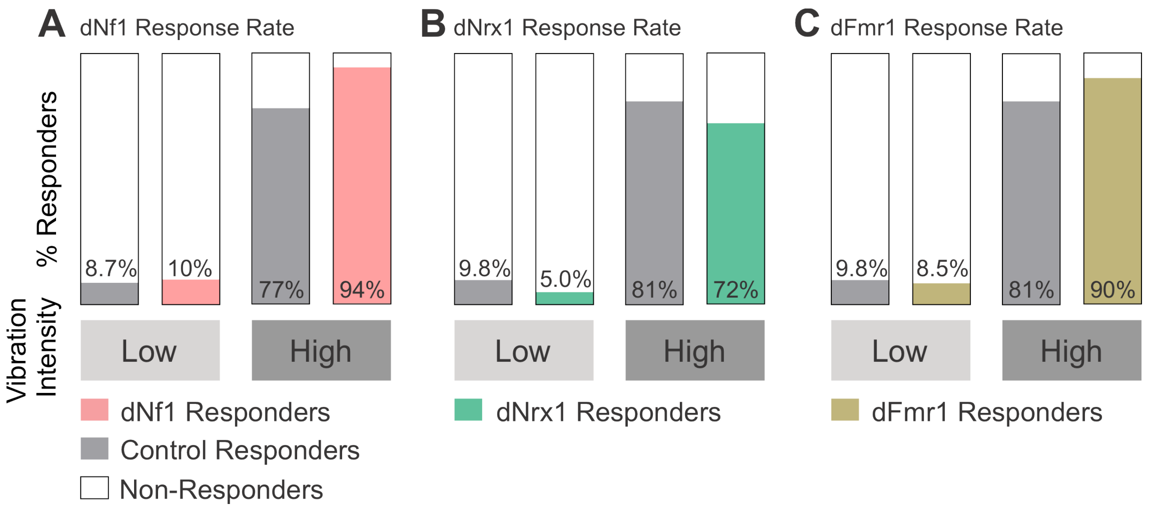


**Supplementary Figure 4. Response rates to low- and high-intensity vibration are comparable between NDD mutants and controls.** Adult female flies were exposed to low-intensity (0.18) or high-intensity (0.31) vibration using a Universal Vibration Stage (UVS-01). Locomotor activity was recorded before and during vibration as described for Fig. 2. Only flies that were awake at vibration onset were included. Flies were classified as responders if their absolute change in activity during the first 5 min of vibration exceeded two standard deviations of their activity during the 15 min immediately preceding vibration onset. Stacked bars show the percentages of responders and non-responders among dNf1 mutants and their genetic controls (A), dNrx1 mutants and their genetic controls (B), and dFmr1 mutants and their genetic controls (C). Colored and gray regions indicate mutant and control responders, respectively; white regions indicate non-responders. Few flies responded at the lower vibration intensity, whereas most flies responded at the higher intensity. At low intensity, sample sizes were: control, n = 23; dNf1, n = 19; shared control for dNrx1 and dFmr1, n = 51; dNrx1, n = 40; and dFmr1, n = 47. At high intensity, sample sizes were: control, n = 27; dNf1, n = 17; shared control for dNrx1 and dFmr1, n = 36; dNrx1, n = 31; and dFmr1, n = 30. Within each intensity, responder proportions were compared between mutants and their corresponding controls using two-sided Fisher's exact tests. No mutant-control comparison was statistically significant (all $P > 0.05$).

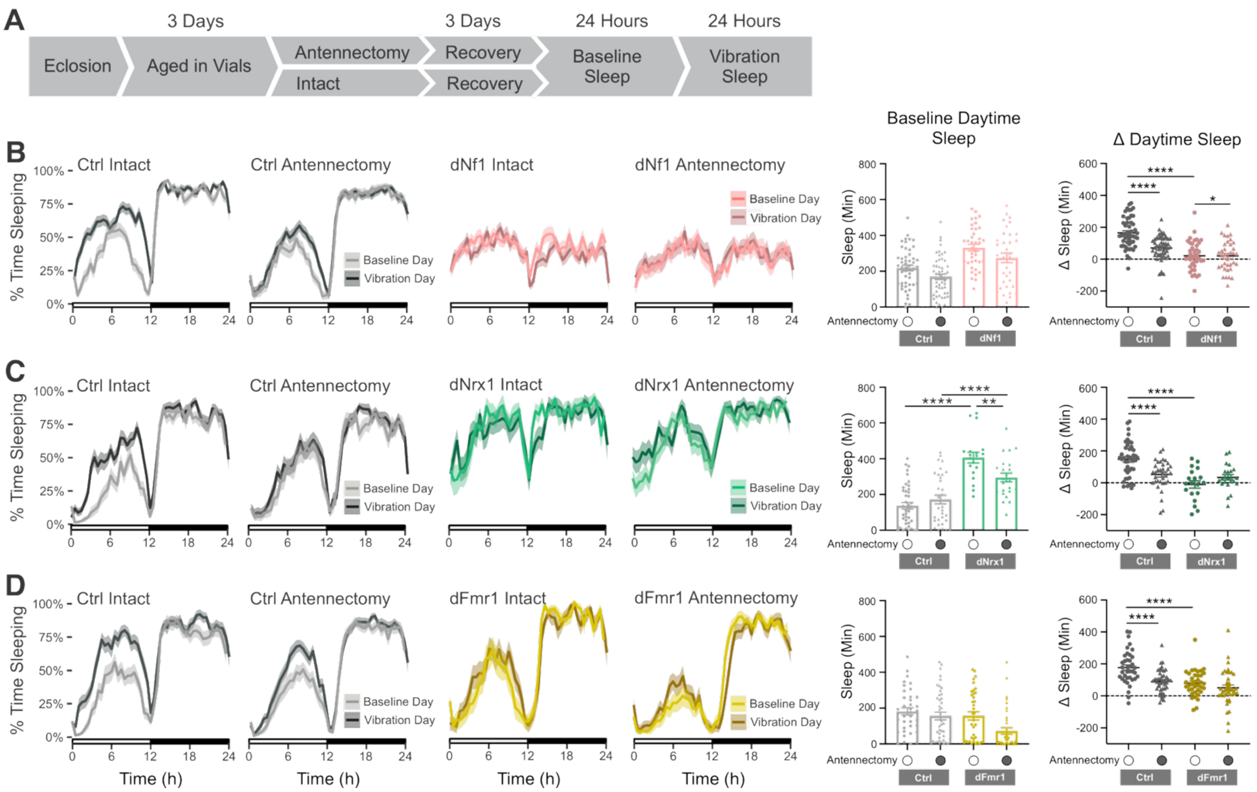


**Supplementary Figure 5. Antennectomy reduces vibration-induced sleep in controls but does not restore it in NDD mutants.** (A) Experimental design. Following eclosion, adult female flies were aged in vials for 3 days. The antennae were then surgically removed or left intact, and flies were allowed to recover for 3 days. Sleep was subsequently recorded for 24 h under baseline conditions and for 24 h during continuous gentle vibration. (B–D) Averaged 24-h sleep traces during the baseline and vibration days for intact and antennectomized flies in dNf1 (B), dNrx1 (C), and dFmr1 (D) mutants and their corresponding genetic controls. Lines show the mean, and shaded regions indicate SEM. Middle graphs show baseline daytime sleep (ZT0–12). Right graphs show vibration-induced sleep, calculated for each fly as the change in daytime sleep between the vibration and baseline days (Δ daytime sleep = vibration-day sleep − baseline-day sleep, ZT0–12). Antennectomy reduced vibration-induced sleep in control flies but did not impact vibration-induced sleep in any of the mutant strains. Points represent individual flies; bars show mean ± SEM. Sample sizes for intact and antennectomized flies, respectively, were: control, n = 50 and 53, and dNf1, n = 39 and 35 (B); control, n = 42 and 34, and dNrx1, n = 19 and 23 (C); and control, n = 38 and 43, and dFmr1, n = 37 and 40 (D). Baseline daytime sleep and vibration-induced sleep were analyzed separately for each mutant–control comparison using ordinary two-way ANOVA, with genotype and antennectomy status as between-subject factors. The genotype-by-antennectomy interaction for baseline daytime sleep was significant only in the dNrx1 comparison, whereas the interaction for vibration-induced sleep was significant in all three mutant–control comparisons. When the interaction was significant, uncorrected Fisher's least significant difference tests were used for multiple comparisons. *$P < 0.05$; **$P < 0.01$; ***$P < 0.001$; ****$P < 0.0001$.

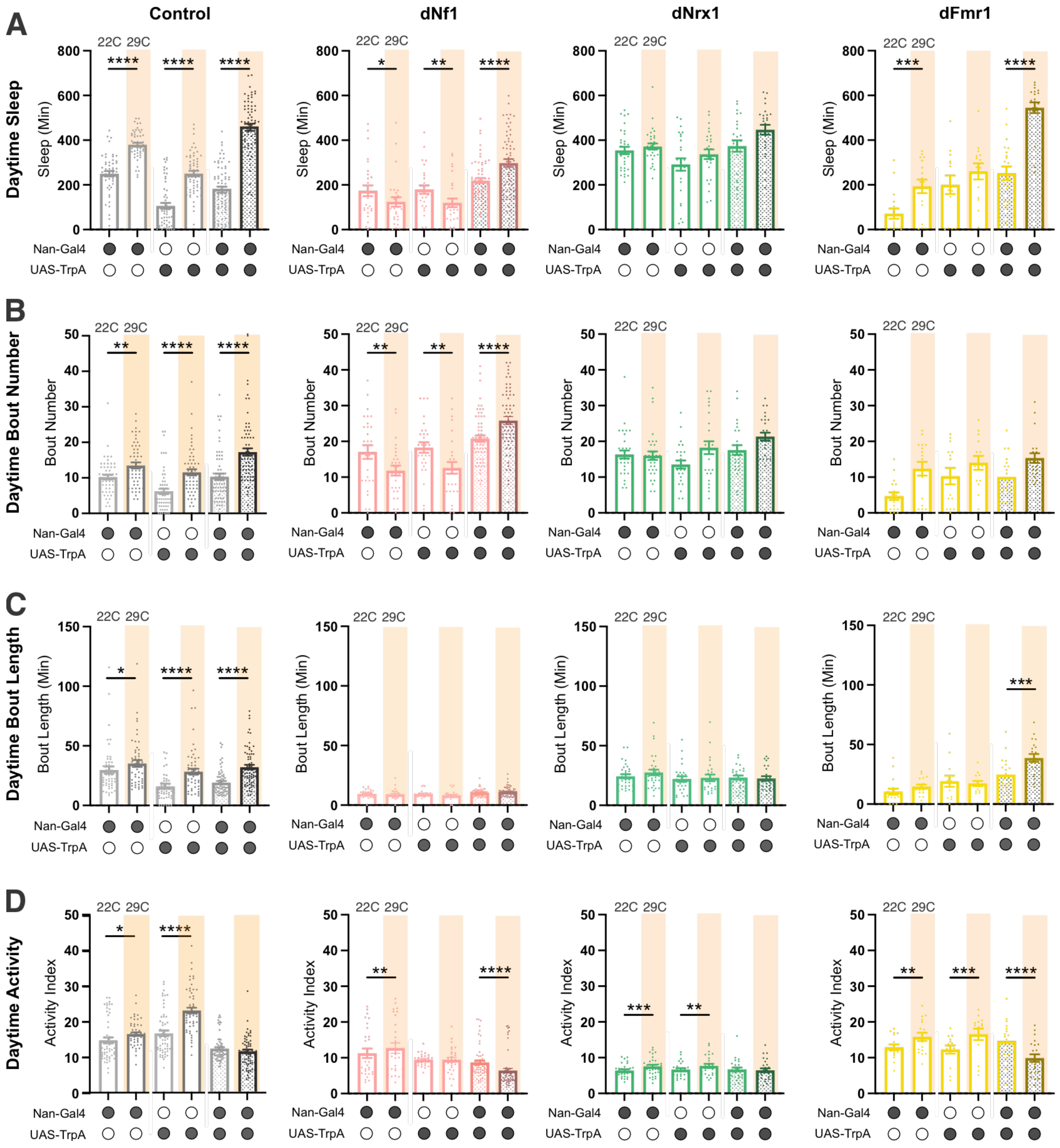


**Supplementary Figure 6. Effects of thermogenetic Nan+ neuron activation on daytime sleep architecture and waking activity.**
Raw daytime sleep parameters were measured at 22°C and 29°C in flies carrying *Nan-GAL4* alone, *UAS-TrpA1* alone, or both transgenes (*Nan-GAL4 > UAS-TrpA1*) in control, *dNf1*, *dNrx1*, and *dFmr1* genetic backgrounds. Temperature was increased from 22°C to 29°C to activate TrpA1 in Nan+ neurons. Orange shading denotes measurements collected at 29°C. (A) Total daytime sleep. (B) Daytime sleep-bout number. (C) Mean daytime sleep-bout length. *(*D) Daytime waking activity index. Each point represents an individual fly; bars show mean ± SEM. Measurements at 22°C and 29°C were obtained from the same flies. Sample

sizes for Nan-GAL4/+, UAS-TrpA1/+, and Nan-GAL4 > UAS-TrpA1, respectively, were: control, n = 53, 58, and 67; dNf1, n = 28, 28, and 62; dNrx1, n = 32, 26, and 26; and dFmr1, n = 17, 14, and 21. Within each genetic background, each parameter was analyzed using a two-way mixed-design repeated-measures ANOVA, with transgene genotype as the between-subject factor and temperature as the within-subject factor. When the genotype-by-temperature interaction was significant, Holm–Šídák multiple-comparisons tests were used to compare 22°C and 29°C within each transgene genotype; significant comparisons are indicated above the corresponding bars. *$P < 0.05$; **$P < 0.01$, ***$P < 0.001$; ****$P < 0.0001$.

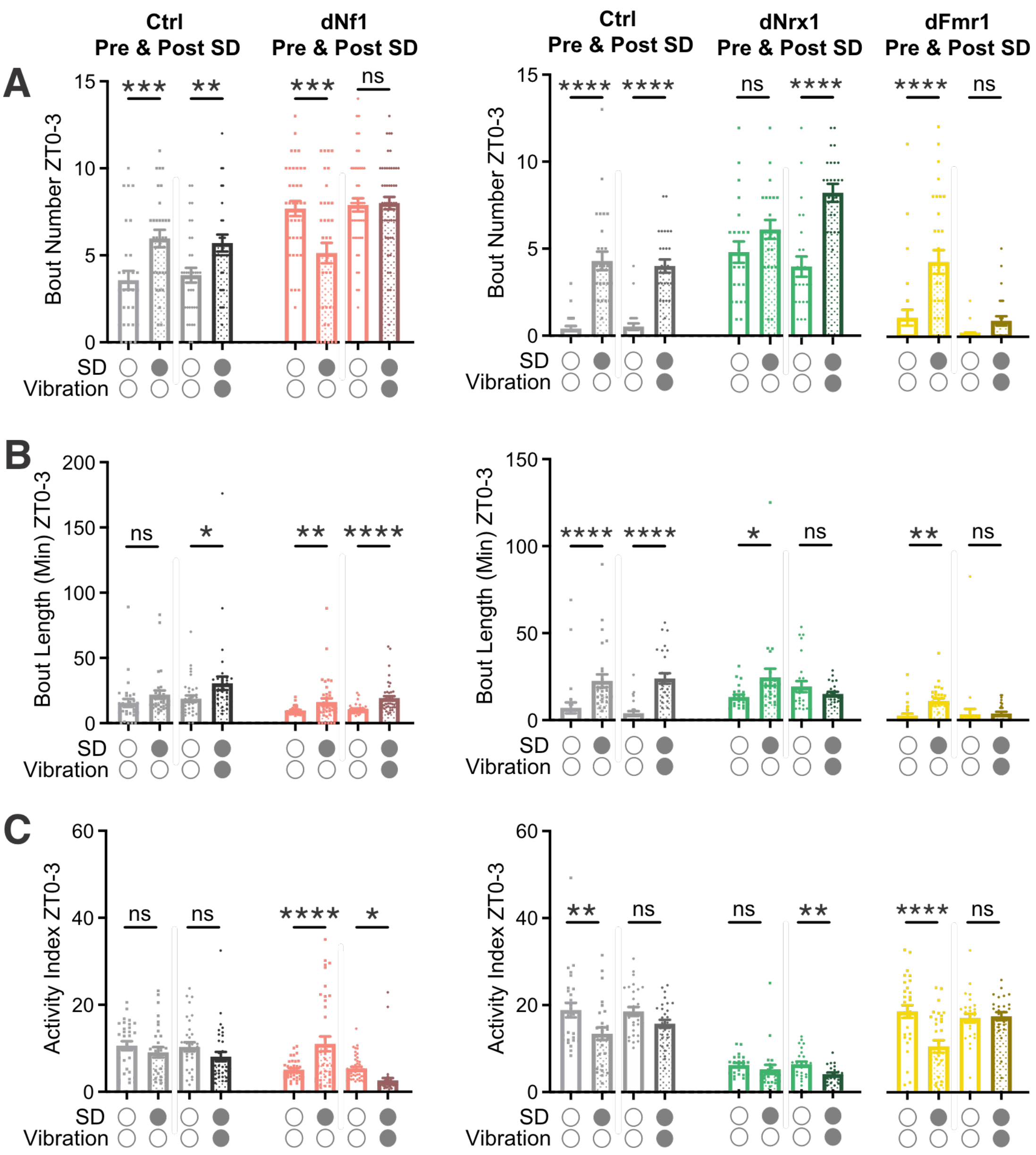

**Supplementary Figure 7. Vibration differentially alters sleep architecture and waking activity following sleep deprivation.** Raw sleep-bout number (A), mean sleep-bout length (B), and waking activity index (C) were quantified during the ZT0–ZT3 interval under baseline conditions and following sleep deprivation (SD), with recovery occurring under either sham or gentle-vibration conditions. Points represent individual flies; bars show mean ± SEM. Sample sizes were as follows: genetic control for dNf1, sham n = 42 and vibration n = 39; dNf1, sham n = 38 and vibration n = 49; shared genetic control for dNrx1 and dFmr1, sham n = 27 and vibration n = 29; dNrx1, sham n = 23 and vibration n = 25; and dFmr1, sham n = 30 and vibration n = 28. Each parameter was analyzed separately within each genotype using a two-way mixed-design ANOVA, with recording condition (baseline vs. post-SD) as a within-subject factor and rebound treatment (sham vs. vibration) as a between-subject factor. The condition-by-treatment interaction was not significant for any parameter in either control background. Significant interactions were detected for all three parameters in dNf1 and dFmr1 mutants and for bout number and mean bout length in dNrx1 mutants. Holm–Šídák multiple comparisons tests shown above the graphs compare baseline and post-SD values within each recovery-treatment group. ns, $P > 0.05$; *$P < 0.05$; **$P < 0.01$; ***$P < 0.001$; ****$P < 0.0001$.
